# AAV-mediated generation of neurons in injury-induced and inherited retinal degenerations

**DOI:** 10.1101/2025.11.28.688728

**Authors:** Mengling Yang, Sherine Awad, Amelia Chung, Jenna Dang, Madelyn Chong, Zachary Flickinger, Kwoon Y. Wong, Thanh Hoang

## Abstract

Loss of retinal neurons is a leading cause of irreversible vision impairment, yet adult mammalian retinas lack the ability to regenerate these cells. In this study, we developed an adeno-associated virus (AAV)-based strategy to reprogram endogenous Müller glia into retinal neurons. Using stringent genetic lineage tracing, immunohistochemistry, electrophysiology and single-cell multiomic profiling, we show that AAV delivery of a stabilized, phospho-insensitive Neurogenin2 variant (Neurog2-9SA) efficiently converts Müller glia into multiple types of retinal neurons, including bipolar, starburst amacrine, and a small population of photoreceptor-like cells, in both injury-induced and inherited retinal degeneration models. The generated neurons exhibit electrical properties of retinal neurons, light-evoked responses and integrate into existing retinal circuitry. Single-cell multiomics analysis reveal that Neurog2-9SA induces reprogramming by remodeling chromatin accessibility, activating neurogenic transcriptional networks, and represses glial identity programs. Inhibiting Notch signaling markedly enhances reprogramming efficiency. Together, these findings establish Neurog2-9SA as a potent and clinically relevant factor for AAV-mediated reprogramming and provide a foundation for approaches to regenerate neuronal cells and restore function in retinal diseases.

## Introduction

Neurons in the retina, like those throughout the central nervous system, are highly susceptible to injury and disease. The loss of retinal neurons underlies major retinal diseases, including retinitis pigmentosa, age-related macular degeneration, diabetic retinopathy and glaucoma ^1^. Current treatments can alleviate symptoms or slow disease progression, but none can restore neurons that have been lost. Cell transplantation approaches are under clinical evaluation, but the efficacy, long-term survival of transplanted cells, immunogenicity and safety of such therapies remain uncertain ^2,3^. Gene therapies targeting individual disease-causing mutations hold promise for treating inherited retinal dystrophies ^4^. However, there are multiple challenges to this approach, including a large number of genes associated with inherited retinal dystrophies, large coding sequences incompatible with standard AAV vectors, cases with unknown or polygenic diseases and disease conditions. Therefore, a broadly effective strategy capable of addressing diverse genetic causes and disease conditions is highly desirable.

One promising strategy is to harness the regenerative potential of Müller glia cells, the principal glial cells of the retina. In non-mammalian vertebrates, such as zebrafish and Xenopus, Müller glia possess a remarkable capacity to re-enter a progenitor-like state after injury and regenerate all major retinal neuron classes ^2,5^. By contrast, Müller glia in adult mammals cannot spontaneously regenerate neurons. Recent advances in reprogramming mammalian Müller glia into retinal neurons have opened new avenues for regenerative therapies. We have previously shown that disrupting key transcriptional programs by deleting NFIs transcription factors ^6^ or inhibiting Notch signaling ^7,8^, induces Müller glia to reprogram into retinal interneurons and a small number of cells expressing rod photoreceptor markers. Additionally, other studies have shown that transgenic overexpression of the transcription factor Ascl1 in combination with chemical treatments induces Müller glia reprogramming into retinal interneurons and ganglion-like cells ^9,10^. Despite these advances, these strategies generally rely on complex transgenic mouse models and combinatorial treatments, limiting their translational potential.

Adeno-associated virus (AAV) vectors offer a clinically relevant platform for *in vivo* reprogramming. However, previous AAV-based attempts ^11–16^ have been complicated by leaky GFAP promoter-driven expression of the reporter in endogenous neurons, leading to misinterpretation of lineage outcomes^17–22^. To enable translational progress, there is a critical need for an AAV-based reprogramming platform that is genetically rigorous, mechanistically well-defined and provides lineage-specific results.

Using both injury-induced and inherited degeneration models, we sought to determine if the proneural bHLH transcription factor Neurogenin 2 (Neurog2) could reprogram mammalian Müller glia and regenerate retinal neurons *in vivo*. We show that AAV-mediated expression of a stabilized, phospho-insensitive Neurog2 variant (Neurog2-9SA), but not wild-type Neurog2, efficiently converts adult Müller glia into multiple classes of retinal neurons. Furthermore, the generated neurons exhibit molecular profiles, morphologies and light-evoked electrophysiological responses comparable to native retinal neurons. Single-cell multiomics reveals that Neurog2-9SA remodels chromatin accessibility, activates neurogenic transcriptional programs and suppresses Müller glia identity programs. Finally, we show that inhibition of Notch signalling enhances reprogramming efficiency, providing a synergistic strategy for boosting neuronal regeneration. Our findings establish Neurog2-9SA as a promising candidate for clinically relevant strategies to regenerate retinal neurons..

## Results

### 1. AAV-mediated expression of phospho-insensitive Neurog2 induces Müller glia reprogramming and neuronal generation after injury

Neurog2 plays a central role in retinal neurogenesis. Neurog2-positive retinal progenitors generate all major classes of retinal neurons ^23^, including photoreceptors and bipolar cells ^24,25^. To define the temporal dynamics of Neurog2 expression, we re-analyzed our previously published single-cell RNA sequencing (scRNA-seq) datasets from developing mice and humans^26,27^ and determined that Neurog2 expression is enriched in the neurogenic retinal progenitors that give rise to all major retinal neuron types (Fig.S1A). Consistent with a potential role in glial reprogramming, Neurog2 expression is re-activated during Müller glia reprogramming when Nfi transcription factors and Notch signaling are deleted ^7^ (Fig.S1B). Although Neurog2 overexpression has been reported to induce limited reprogramming of neonatal Müller glial cells *in vitro* ^28^, it is unknown if Neurog2 can reprogram adult Müller glial cells *in vivo*.

To test this, we delivered Cre-inducible FLEX AAV2/7m8 vectors encoding either mScarlet3 alone, wild-type Neurog2, or a variant form of Neurog2, in which 9 Serine sites in flexible regions are changed to Alanine (thereafter Neurog2-9SA) which prevents phosphorylation and enhances Neurog2 protein stability ^29,30^, into adult GlastCreERT/Sun1GFP mice (Fig.1A). The FLEX AAV system was used to minimize leaky expression in native neurons, which has been observed with direct GFAP promoter-driven AAVs ^19^. Mice received five daily intraperitoneal injections of Tamoxifen to simultaneously induce GFP, Neurog2 and mScarlet3 expression specifically in Müller glial cells (Fig.1B). All 3 AAV constructs efficiently transduce Müller glial cells with 60-70% of GFP+ cells co-expressing mScarlet3 (Fig.1C, E). As expected, Neurog2-9SA generated higher protein levels than wild-type Neurog2 (Fig.S1C).

**Figure 1.**
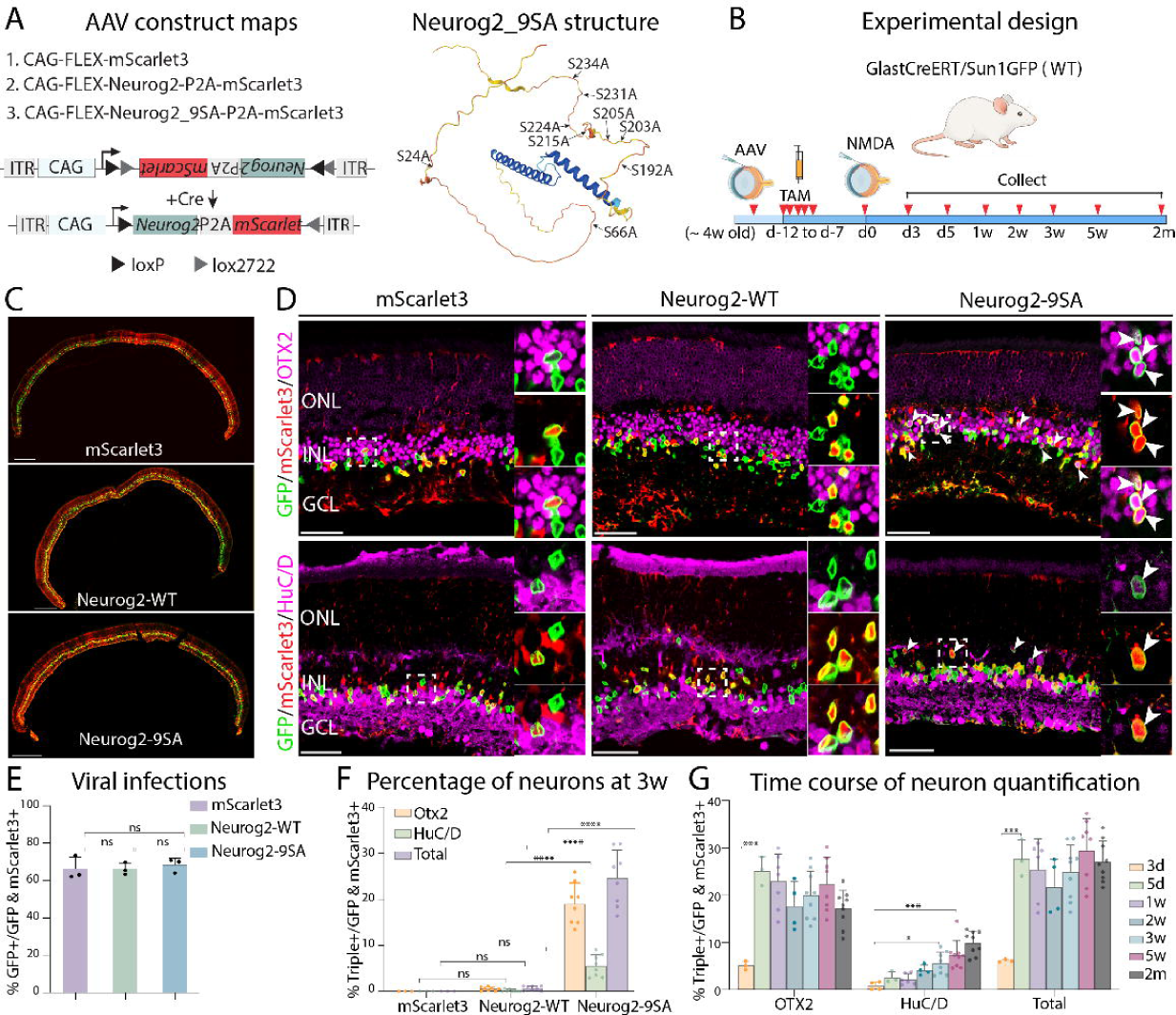
AAV-mediated expression of stable form of Neurog2 induces Müller glia reprogramming to generate retinal neurons. (A). Left: Schematic maps of three FLEX AAVs: CAG-FLEX-mScarlet3 (reporter control), CAG-FLEX-Neurog2-P2A-mScarlet3 (Neurog2-WT), and CAG-FLEX-Neurog2_9SA-P2A-mScarlet3 (Neurog2-9SA). Right, alpha-fold structure of Neurog2_9SA carrying 9 serine-to-alanine substitutions (9SA).(B) Experimental design and sample collection. (C, E) Representative retinal wholemount immunostaining for GFP (Müller glial lineage) and mScarlet3 (AAV reporter), and quantification of Müller glia-targeted transduction (fraction of mScarlet3⁺ cells that are GFP⁺) across mScarlet3, Neurog2-WT and Neurog2-9SA groups at 2 weeks post tamoxifen injection. Scale bars, 300 µm. (D) Representative immunostained images of GFP (green), mScarlet3 (red), and either Otx2 or HuC/D (magenta) at 3 weeks post NMDA injury. White arrowheads mark triple-positive GFP⁺/mScarlet3⁺/marker⁺ Müller glia-derived neurons. These generated neurons often show weaker GFP intensity than typical Müller glia. Insets show higher-magnification examples. Scale bars, 50 µm. (F) Quantification of percentage (mean ± SD) of GFP⁺/mScarlet3⁺ cells that express Otx2 or HuC/D in each group; points represent individual retinas. (G) Quantification of generated neurons in Neurog2-9SA-overexpressed retinas over serial time points. Proportion (mean ± SD; points = individual retinas) of GFP⁺/mScarlet3⁺ cells that are Otx2⁺ or HuC/D⁺ at 3 d, 5 d, 1 w, 2 w, 3 w, 5 w, and 2 m post-injury, with the total triple-positive fraction shown for reference. Error bars indicates mean standard deviation. Significance was determined via one-way ANOVA with Tukey’s multiple comparison test: *p < 0.05, **p < 0.01, ***P < 0.001, ****p < 0.0001. TAM: Tamoxifen; ONL: outer nuclei layer; INL: inner nuclear layer; GCL: ganglion cell layers.

Following NMDA-induced retinal injury, a paradigm that activates Müller glia but does not induce spontaneous regeneration ^6,31^, only Neurog2-9SA induced a substantial reprogramming of Müller glia into retinal neurons (Fig.1D, F). In contrast to the mScarlet3 control or wild-type Neurog2, Neurog2-9SA-infected Müller glia gave rise to OTX2⁺ neurons (∼20%) and HuC/D⁺ neurons (∼6%). Time-course analysis revealed distinct temporal dynamics for neurogenesis (Fig.1B, G). We found that OTX2+ neurons derived from Müller glia rapidly emerged as early as 3 days post-injury, peaked at 5 days and remained stable for at least 2 months. Meanwhile, HuC/D+ neurons appeared later and accumulated gradually, peaking at 5 weeks post injury (Fig.1G). Together, these findings demonstrated that AAV-mediated delivery of Neurog2-9SA, but not wild-type Neurog2, is sufficient to reprogram Müller glia and generate retinal neurons in the adult mice following injury.

### 2. Neurog2-9SA overexpression reprograms Müller glia to generate different types of retinal neurons

To further characterize the molecular features of Müller glia-derived neurons, we performed scRNA-Seq on FACS-purified GFP+ cells from mScarlet3 control and Neurog2-9SA-transduced retinas at 5 and 8 weeks after NMDA injury (Fig.2A). After quality-control filtering, we obtained 9,701 cells from mScarlet3 controls and 27,732 and 11,486 cells from Neurog2-9SA at 5 and 8 weeks respectively (Fig. 2B). Unbiased UMAP clustering combined with established retinal cell markers were used to annotate cell identities ^26^. As expected, GFP+ cells from mScarlet3 control group consisted primarily of Müller glial and a small fraction of retinal neurons, likely contaminants introduced during FACS isolation (Fig.2C, D).

**Figure 2.**
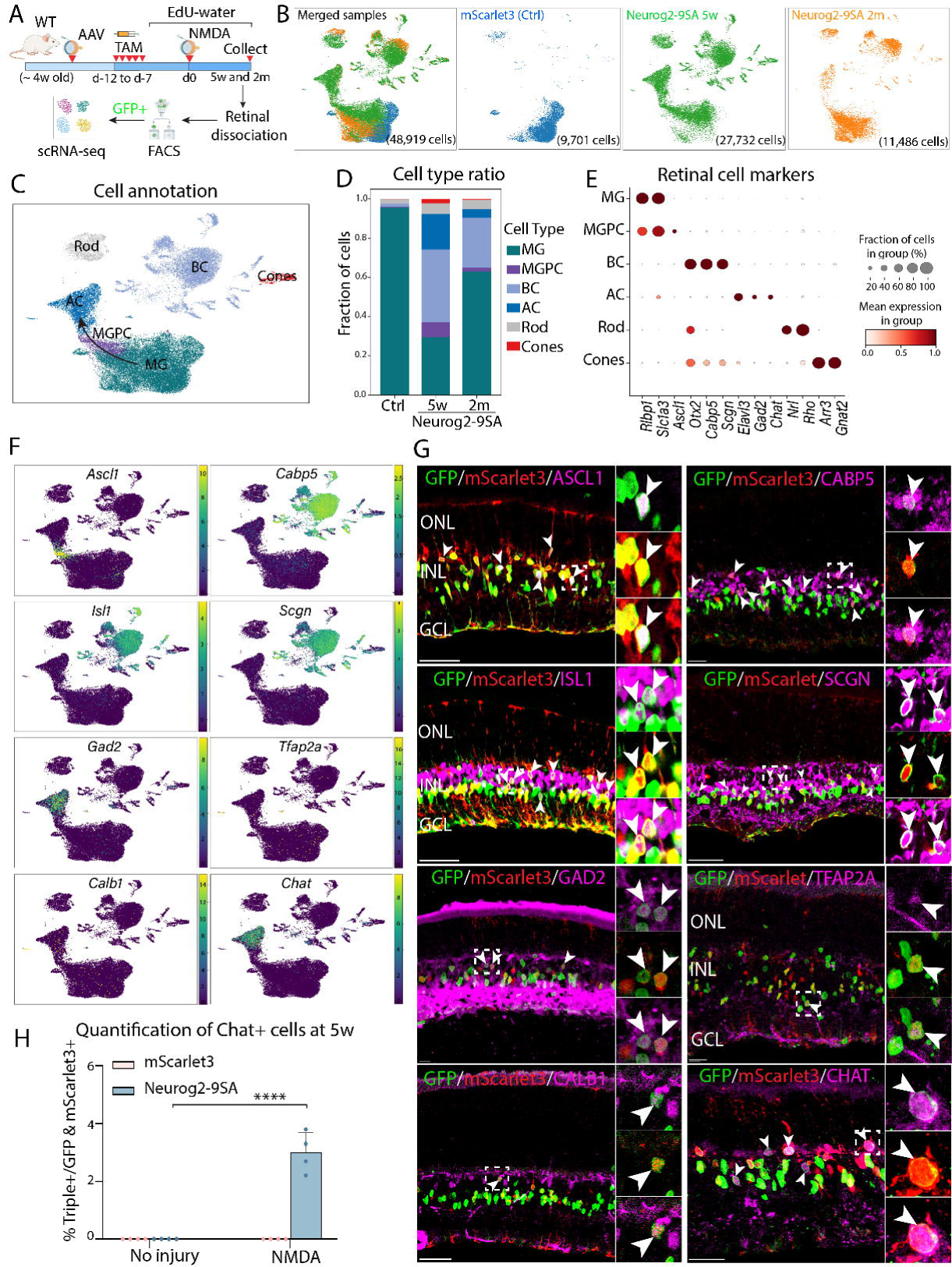
Molecular characterization of neurons generated by Neurog2-9SA expression. (A) Experimental workflow for purified GFP+ cells and scRNA-seq. (B) UMAPs of both integrated and individual datasets showing control (mScarlet3) and Neurog2-9SA samples at 5 weeks and 2 months after NMDA injury. Numbers on panels indicate the number of high-quality cells for each sample. (C,D) Cell-type annotation and proportion of cell types across conditions. Neurog2-9SA samples show a reduced Müller glia fraction with a corresponding increase in retinal neurons compared with the control group. (E) Dot plot of selected markers showing downregulation of resting Müller glial genes (Rlbp1 and Slc1a3) and upregulation of neurogenic (Ascl1) and neuronal markers. (F,G) Feature plots showing expression of bHLH neurogenic transcription factor Ascl1, cone bipolar markers (Cabp5, Isl1 and Scgn), amacrine markers (Gad2,Tfap2a, Calb1 and Chat). Representative immunostained images validate the scRNA-seq data. White arrowheads indicate triple-positive GFP⁺/mScarlet3⁺/marker⁺ cells; Müller glia-derived neuron-like cells typically show lower GFP intensity than Müller glia. (H) Quantification of CHAT⁺ Müller glia-derived neurons (mean ± SD) among GFP⁺/mScarlet3⁺ cells in mScarlet3 control versus Neurog2-9SA retinas, with and without NMDA injury. Dot points represent individual retinas. Error bars indicates ± mean standard deviation. Significance was determined via one-way ANOVA with Tukey’s multiple comparison test: *p < 0.05, **p < 0.01, ***p < 0.001, ****p < 0.0001. MG, Müller glia, MGPC; Müller glia-derived progenitors; BC, bipolar cell; AC, amacrine cell; ONL: outer nuclei layer; INL: inner nuclear layer; GCL: ganglion cell layers Scale bars, 50 µm.

In contrast, Neurog2-9SA-overexpression drove extensive Müller glia reprogramming, yielding multiple distinct cell populations, including Müller glia-derived progenitor cells (MGPCs) that transition into amacrine-like cells and bipolar-like cells (Fig.2C, D, Fig.S2A). Neurog2-9SA expression resulted in the de-differentiation of Müller glia, as evidenced by the downregulation of Müller glial markers, such as *Rbp1*, *Slc1a3* and *Lhx2* (Fig.2E, Fig.S2B,C, Table 1). We observed that MGPCs expressed the bHLH neurogenic transcription factor, *Ascl1*, which we confirmed by immunohistochemistry (Fig.2E-G).

The scRNA-Seq analysis shows that the bipolar-like cell cluster was well separated from the MGPC cluster in UMAP space (Fig.2C), suggesting that the regeneration of bipolar cells is largely completed at this timepoint. This result is consistent with our serial time-course immunostaining data (Fig.1G). The bipolar-like cells expressed the pan-bipolar cell marker *Otx2* (Fig.2E, Fig.S2A) along with cone bipolar subtype-associated markers *Cabp5*, *Isl1* and *Scgn* (Fig.2E-G, Table 1). Amacrine-like cells expressed HuC/D and *Gad2*, with smaller subsets expressing *Tfap2a* and *Calb1* (Fig.2F, G). Interestingly, we identified a population of amacrine-like cells expressing ChAT, the vesicular acetylcholine transporter, VAChT (*Slc18a3*) and *Megf11*, indicating the generation of starburst amacrine cells (Fig.2F,G, Fig.S2A, Table 1). Immunostaining data showed that ∼3% of AAV-infected GFP+ cells were ChAT+ starburst amacrine-like cells (Fig.2H).

We also assessed the generation of other retinal cell types. We observed a small number of GFP+ generated neurons expressing rod photoreceptor marker NRL (Fig.S2D), but we did not detect obvious generation of cones (ARR3), rod bipolar cells (PRKCA) or retinal ganglion cells (RGC) (ATOH7, BRN3A and RBPMS) (Fig.S2D).

We next asked whether Neurog2-9SA overexpression induces Müller glia proliferation during reprogramming. To label proliferating cells, mice were injected with Neurog2-9SA AAV and treated with EdU in the drinking water (Fig.2A). NMDA injury induced minimal Müller glia proliferation, and we observed no significant difference in the number of EdU+/GFP+ cells between Neurog2-9SA and mScarlet3 controls (Fig.S2E). Consistent with this, our scRNA-Seq data did not detect obvious expression proliferation markers *Ki67* and *Cdk1* (Table 1). Together, these data demonstrate that Neurog2-9SA overexpression reprograms Müller glia into multiple neuronal lineages, including bipolar cells, starburst amacrine cells, and a small number of rod photoreceptors, mainly through direct transdifferentiation rather than through a proliferation-driven mechanism.

### 3. Generated neurons morphologically and electrophysiologically resemble retinal neurons

We next assessed the morphological features of neurons generated from Neurog2-9SA-overexpressing Müller glia. In control retinas, immunostaining of mScarlet3 reporter showed that GFP+ Müller glia exhibit typical radial morphology, spanning the outer, inner and ganglion cell layers (Fig.3A). In contrast, GFP+ Müller glia-derived neurons adopt morphologies and axonal/dendritic lamination patterns typical of retinal neurons (Fig.3A). Co-immunostating with retinal markers confirm the identities of these generated neurons as Otx2+ bipolar and HuC/D+ amacrine cells, including ChAT+ starburst amacrine cells.

**Figure 3.**
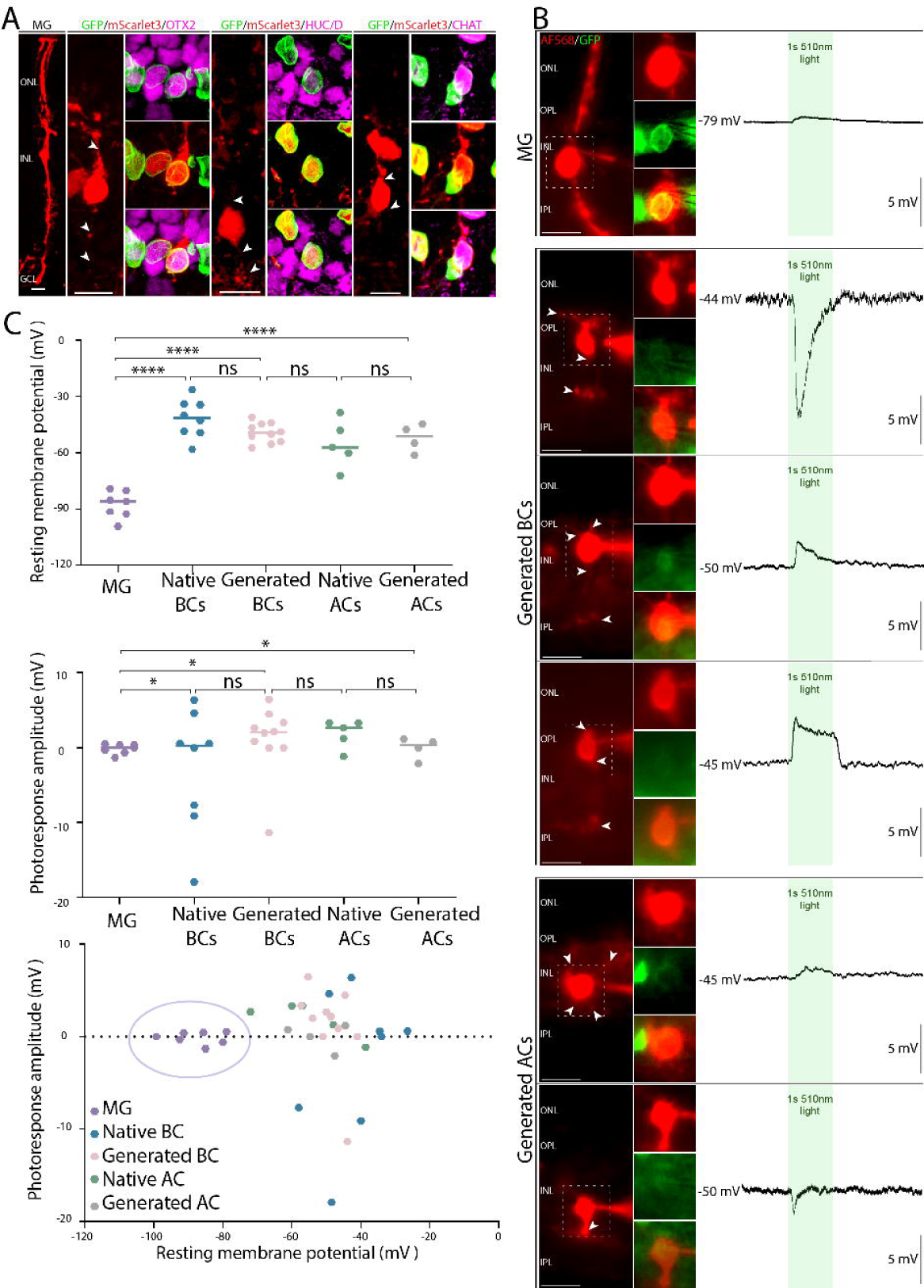
Morphological and electrophysiological characterization of Müller glia-derived neurons. (A) Representative images of morphological characterization of Müller glia and Müller glia-derived neurons with Neurog2-9SA overexpression. Neuronal identities were confirmed by co-immunostaining with neuronal markers OTX2, HuC/D and CHAT. White arrows point to projections of the neuron-like cells. Scale bars, 10 μm. (B) Morphological and electrophysiological analysis of GFP+/mScarlet3+ Müller glia and Müller glia-derived bipolar and amacine-like cells. Cells were filled with Alexa568 dye during the whole-cell recording process to enhance visualization of cell morphology. Müller glia-derived neurons exhibited higher resting membrane potential than typical Müller glial cells, and responded to light stimulation. (C) Quantifications of membrane potential and photoresponse amplitude for Müller glia (n=7), Müller glia-derived neurons (n=14) and native bipolar (n=8) and native amacrine neurons (n=5). Generated bipolar and amacrine cells exhibited similar membrane potential and photoresponse amplitude with native retinal neurons. Dot points represent individual cells. Error bars indicates mean ± SD. Significance was assessed using T-tests and one-way ANOVA, with analyses conducted on absolute values. *P < 0.05, ***P < 0.001. GCL gangling, INL inner nuclear layer, ONL outer nuclear layer. : *P < 0.05, **p < 0.01, ***p < 0.001, ****p < 0.0001. ONL: outer nuclei layer; OPL: outer plexiform layer; IPL: inner plexiform layer; INL: inner nuclear layer; GCL: ganglion cell layers.

To determine whether generated neurons acquire functional electrophysiological properties, we performed whole-cell patch clamp recording from individual GFP+/mScarlet3+ cells in retinal slices between 5 weeks to 8 weeks after NMDA injury. Alexa Fluor 568 dye was included in the micropipette solution to enhance visualization of cell morphologies. As expected, GFP+ Müller glia exhibited very negative resting membrane potentials and only sluggish, low-amplitude light-evoked responses (Fig.3B, C). In contrast, GFP+/mScarlet3+ generated cells with bipolar- or amacrine-like morphologies showed significantly more positive resting membrane potentials. Their light-evoked responses were faster and larger than those of the GFP+ Müller glia (Fig.3B,C). Overall, 8 of 10 bipolar-like generated neurons and 3 of 4 amacrine-like generated neurons responded to light, with resting potentials and photoresponse amplitudes comparable to those of native retinal neurons (Fig.3C).

Together, these results demonstrate that neurons generated by Neurog29SA not only adopt appropriate molecular identities and morphologies but also develop functional light-evoked activity. Their ability to receive photoreceptor-driven input demonstrates successful integration into existing synaptic circuits.

### 4. Single-cell multiomic analysis reveals the mechanism by which Neurog2 induces neurogenesis

To investigate how Neurog2-9SA induces reprogramming in Müller glia, we performed combined single-nucleus RNA-Seq and ATAC-seq on FACS-purified GFP+ cells from both mScarlet3 control and Neurog2-9SA-treated retinas at 1 week after NMDA injury (Fig.4A). This early time point was chosen to capture early changes in transcription and chromatin accessibility induced by Neurog2-9SA. Integrated analysis of combined snRNA/ATAC profiles was used to define cell states and infer lineage trajectories.

**Figure 4.**
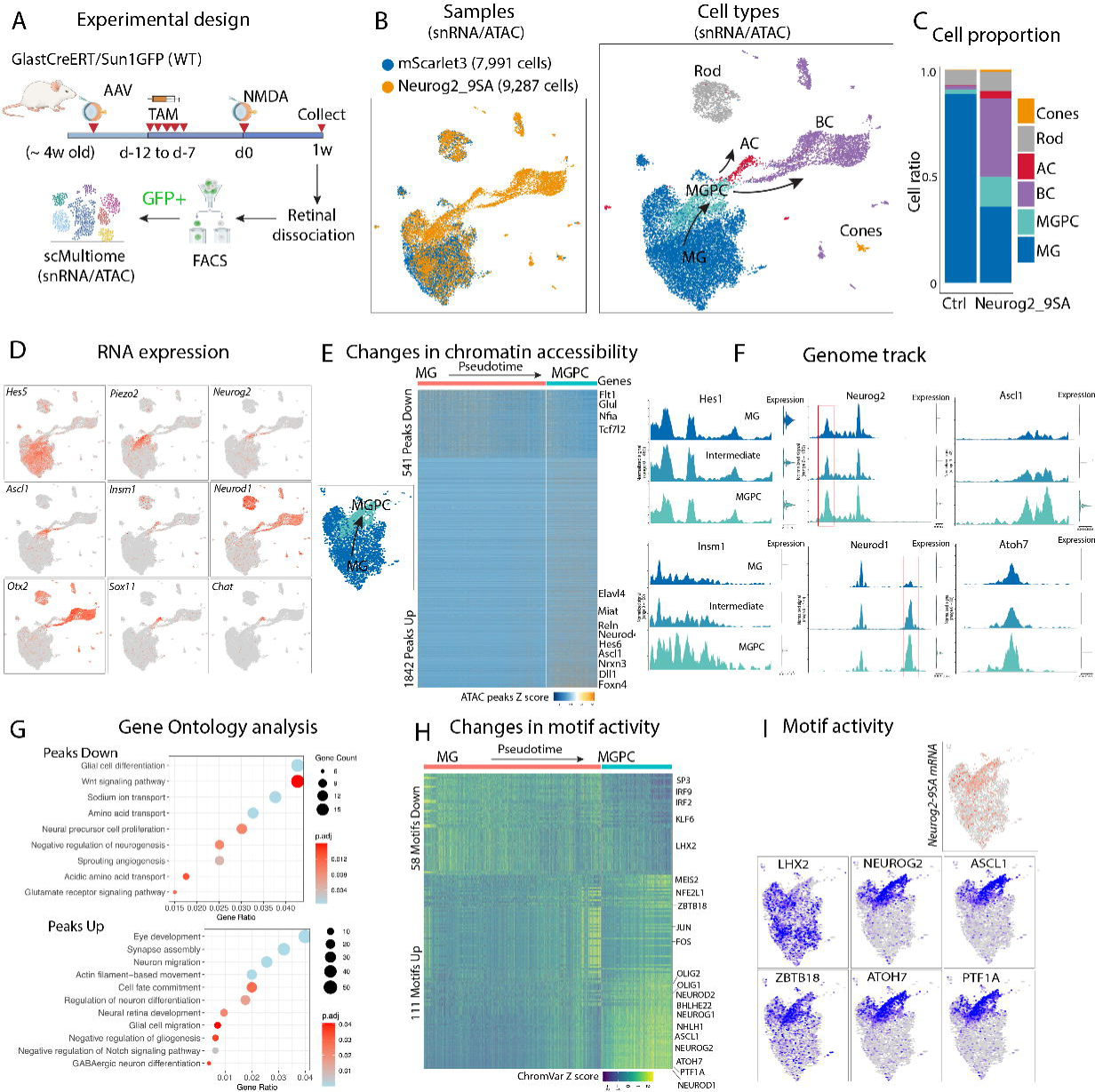
Integrated single-cell Multiomic (snRNA/ATAC) analysis of Müller glia and Müller glia-derived cells following Neurog2-9SA expression. (A) Schematic of experimental design, sample collection and single-cell Multiomic analysis. (B) UMAP plots of scMultiome showing the clustering of the mScarlet3 control and Neurog2-9SA-treated samples, and cell type annotation. (C) Stacked barplot showing the proportion of cells on each cluster across the 2 samples. (D) UMAP plots showing expression of gene markers. (E) Heatmap showing differentially chromatin accessible regions along the pseudotime from MG to MGPC and nearby genes. (F) Genome tracks showing chromatin accessibility changes along the pseudotime from MG to MGPC. (G) Gene ontology of genes near the down and up chromatin accessible regions. (H, I) Heatmap and UMAP plots showing changes in transcription factor motif activity over the pseudotime from MG to MGPC. MG, Müller glia, MGPC; Müller glia-derived progenitors; BC, bipolar cell; AC, amacrine cell.

As expected, we detected the widespread expression of Neurog2-9SA mRNA only in the Neurog2-9SA sample but not in the mScarlet3 control (Fig.S3A). GFP+ cells from the mScarlet3 control sample primarily consisted of Müller glial, with a small fraction of contaminating retinal neurons likely introduced during FACS isolation (Fig.4B, C). In contrast, Neurog2-9SA overexpression produced a clear trajectory from Müller glia cells to MGPCs which subsequently gave rise to bipolar and amacrine cells (Fig.4D, Fig.S3B and Table S2). At this early stage, the majority of the generated neurons displayed bipolar cell identities, expressing markers such as *Otx2*, *Cabp5*, *Scgn* and *Isl1*, while a small population expressed *Sox11* and starburst amacrine cell marker *ChAT* (Fig.4D, Fig.S3B and Table S2). These observations align with our immunostaining data from the same timepoint (Fig.1G).

Pseudotime analyses show that Neurog2-9SA expression induced substantial gene expression changes during the transition from Müller glia to MGPCs. This transition was characterized by the downregulation of glial identity genes, including *Glul*, *Rlbp1*, *Aqp4*, *Hes5* and *Lhx2*, and concurrent upregulation of neurogenic regulator genes, such as *Ascl1*, *Insm1*, *Neurod1/2/4* and endogenous *Neurog2* (Fig.4D, Fig.3C and Table S2). Analysis of chromatin accessibility revealed parallel regulatory changes. Neurog2-9SA expression resulted in 1,842 unregulated and 541 downregulated peaks during the MG-to-MPGC transition, suggesting that Neurog2-9SA broadly opens chromatin accessibility and activates gene expression (Fig.4E, F). Gene Ontology enrichment analysis shows that genes associated down-regulated peaks are involved in glial identify and functions, while genes associated down-regulated peaks are involved in cell migration and neurogenesis (Fig.4G)

Motif enrichment of differentially accessible chromatin regions showed the strong enrichment for Neurog2 motifs, indicating that Neurog2-9SA directly reshapes the chromatin landscape and primes neurogenic programs. Motif activities for transcription factors associated with resting Müller glia, such as *Hes1*, *Lhx2* and *Klf6* were progressively reduced following Neurog2-9SA overexpression, while MGPCs show increased motif activity for neurogenic transcription factors, such as *Ascl1*, *NeuroD1, Zbtb18, Ptf1a* and *Atoh7* (Fig.4H, I). These results suggest that Neurog2-9SA functions as both a transcriptional activator and a pioneer-like factor that opens neurogenic programs while suppressing Müller glial identity.

### 5. Inhibition of Notch signaling enhances Neurog2-9SA-induced neurogenesis

We previously showed that inhibition of Notch signaling through deletion of its core mediator *Rbpj* induces Müller glia de-differentiation and the generation of a limited number of retinal neurons ^7,8^. We also showed that combining Notch inhibition with Nfi*a/b/x* deletion or Oct4 overexpression ^8^ robustly enhances the efficiency of neurogenesis. To test whether Notch inhibition could further augment Neurog2-9SA-mediated reprogramming and regeneration, we performed intravitreal injection of Neurog2-9SA or mScarlet3 control AAVs into Müller glia-specific Rbpj conditional knockout mice (GlastCreERT/Rbpj^flox/flox^/Sun1GFP, hereafter Rbpj cKO). Mice received NMDA injury 1 week after tamoxifen injection or were left uninjured. Retinas were analyzed at 4 and 6 weeks post tamoxifen injections (Fig.5A).

**Figure 5.**
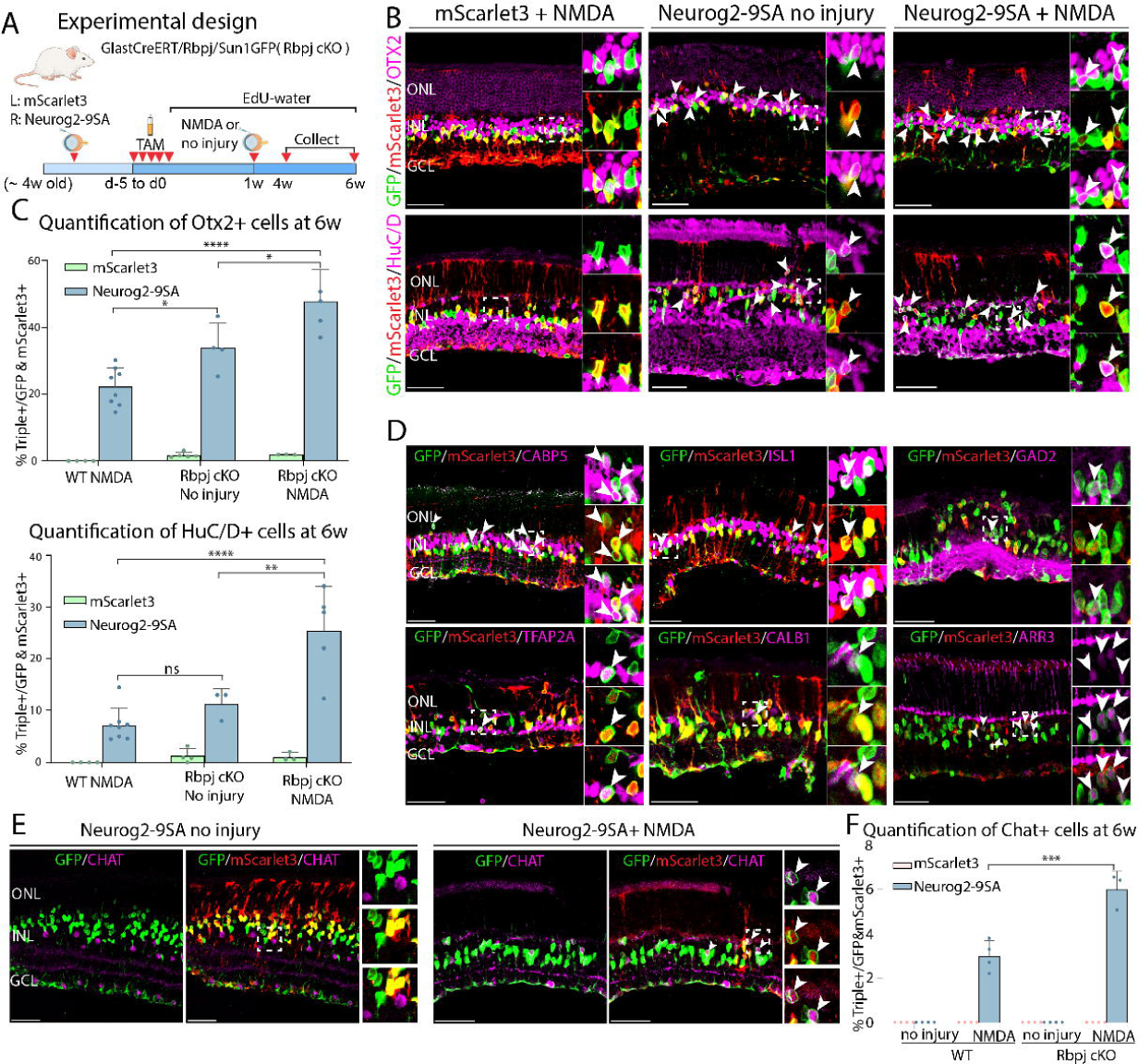
Notch inhibition enhances the efficiency of Neurog2-9SA-induced Müller glia reprogramming. (A) Schematic of experimental design and sample collection. (B) Representative images of immunostaining for GFP (Müller glial lineage), mScarlet3 (AAV reporter), and OTX2 or HuC/D from three Rbpj cKO groups: mScarlet3 + NMDA, Neurog2-9SA without injury, and Neurog2-9SA + NMDA at 5 weeks post NMDA injury (6w post tamoxifen injections). White arrowheads mark triple-positive GFP⁺/mScarlet3⁺/neuronal marker⁺ cells. Müller glia-derived neurons typically display lower GFP intensity than Müller glia. (C) Quantification of Otx2⁺ and HuC/D⁺ percentage among GFP⁺/mScarlet3⁺ cells across three conditions: Rbpj cKO without injury, and injured Rbpj cKO injected with mScarlet3 or Neurog2-9SA, at 5 weeks post NMDA damage (6 weeks post tamoxifen injections). (D) Representative images of Rbpj cKO retinas overexpressing Neurog2-9SA immunolabeled for neuronal markers, including bipolar subtype markers (CABP5 and ISL1), amacrine markers (GAD2,TFAP2A and CALB1) and cone marker (ARR3). (E) Representative images of Rbpj cKO retinas overexpressing Neurog2-9SA immunolabeled for starburst amarine cell marker CHAT at 5w post NMDA injury or 6w post tamoxifen injections (no injury). (F) Quantification of CHAT⁺ Müller glia-derived neurons among GFP⁺/mScarlet3⁺ cells in WT and Rbpj cKO mice injected with mScarlet3 or Neurog2-9SA, with or without NMDA injury at 5 weeks post NMDA injury or 6 weeks post tamoxifen injections (no injury). Error bars indicates mean SD. Significance was determined via one-way ANOVA with Tukey’s multiple comparison test: *p < 0.05, **p < 0.01, ***p < 0.001, ****p < 0.0001. Each dot point represents individual retinas. Arrowheads indicate triple-positive cells. Scale bars, 50 µm.

Neurog2-9SA overexpression in uninjured Rbpj cKO retinas significantly increased the generation of Otx2+ bipolar-like neurons (∼36%) and HuC/D+ amacrine-like neurons (∼12%), relative to both mScarlet3 control and Neurog2-9SA-treated wild-type animals (Fig.5.B, C, Fig.S4D). NMDA injury further enhanced reprogramming efficiency, yielding ∼47% Otx2+ and ∼25% HuC/D+ neurons (Fig.5B, C, FigS4D). Similar to wild-type retinas, immunohistological analysis showed that generated neurons in Rbpj cKO retinas expressed cone bipolar cell subtype markers CABP5, SCGN and ISL1, as well as amacrine markers GAD2, TFAP2A and CALB1 (Fig.5D, Fig.S4A,B).

Notably, Neurog2-9SA overexpression also generated neurons expressing cone photoreceptor markers Arrestin 3 (ARR3) and RXRG in both uninjured and injured Rbpj cKO retinas (Fig.5D, Fig.S4A). We did not observe GFP+ cells expressing rod bipolar marker (PRKCA), additional cone marker (S-Opsin) or RGC markers (BRN3A and RBPMS) (Fig.S4A, B). As observed in injured wildtype retinas, ChAT+ starburst amacrine cells were generated in NMDA-injured Rbpj cKO retinas, but not in uninjured Rbpj cKO retinas, suggesting that injury-induced retinal damage may be required for the regeneration of this highly specialized cell type (Fig.5E, F).

To assess whether Neurog2-9SA overexpression in the Rbpj cKO animals induces Müller glia proliferation during reprogramming, we performed EdU labelling after AAV injection. Only very few EdU+/GFP+ cells were detected in Neurog2-9SA-injected retinas, with no significant difference compared to the mScarlet3 control group (Fig.S5C). These results indicate that, as in the wildtype mice, Neurog2-9SA primarily induces direct transdifferentiation of Rbpj-deficient Müller glia into retinal neurons. Together, these findings demonstrate that inhibition of Notch signaling significantly enhances the reprogramming activity of Neurog2-9SA and expands the neuron type generated from Müller glia.

### 6. Neurog2-9SA overexpression induces the generation of retinal neurons in chronic inherited retinal degeneration

Our data has so far shown that AAV-mediated expression of Neurog2-9SA induces Müller glia reprogramming and generates new neurons in both uninjured and acutely injured retinas. We next asked whether Neurog2-9SA could also induce neurogenesis in a model of chronic retinal degeneration. To address this question, we used the rhodopsin P23H mice, which exhibits progressive photoreceptor degeneration ^32^. We performed intravitreal injections of either Neurog2-9SA or mScarlet3 only AAVs into GlastCreERT/P23H/Sun1GFP mice (hereafter referred to as P23H). Retinas were collected for immunohistological analysis at 4 weeks and 6 weeks after tamoxifen injections (Fig. 6A). Neurog2-9SA overexpression generated Otx2+ bipolar-like and HuC/D+ amacrine-like retinal neurons (Fig. 6B, C), at levels of efficiency comparable to those observed after acute NMDA injury in wild-type mice (Fig.1). The generated neurons expressed bipolar cell markers (CABP5, SCGN and ISL1), amacrine cell markers (CHAT and CALB1), and, at a low frequency, the rod photoreceptor marker NRL (Fig.6E, Fig.S5A).

**Figure 6.**
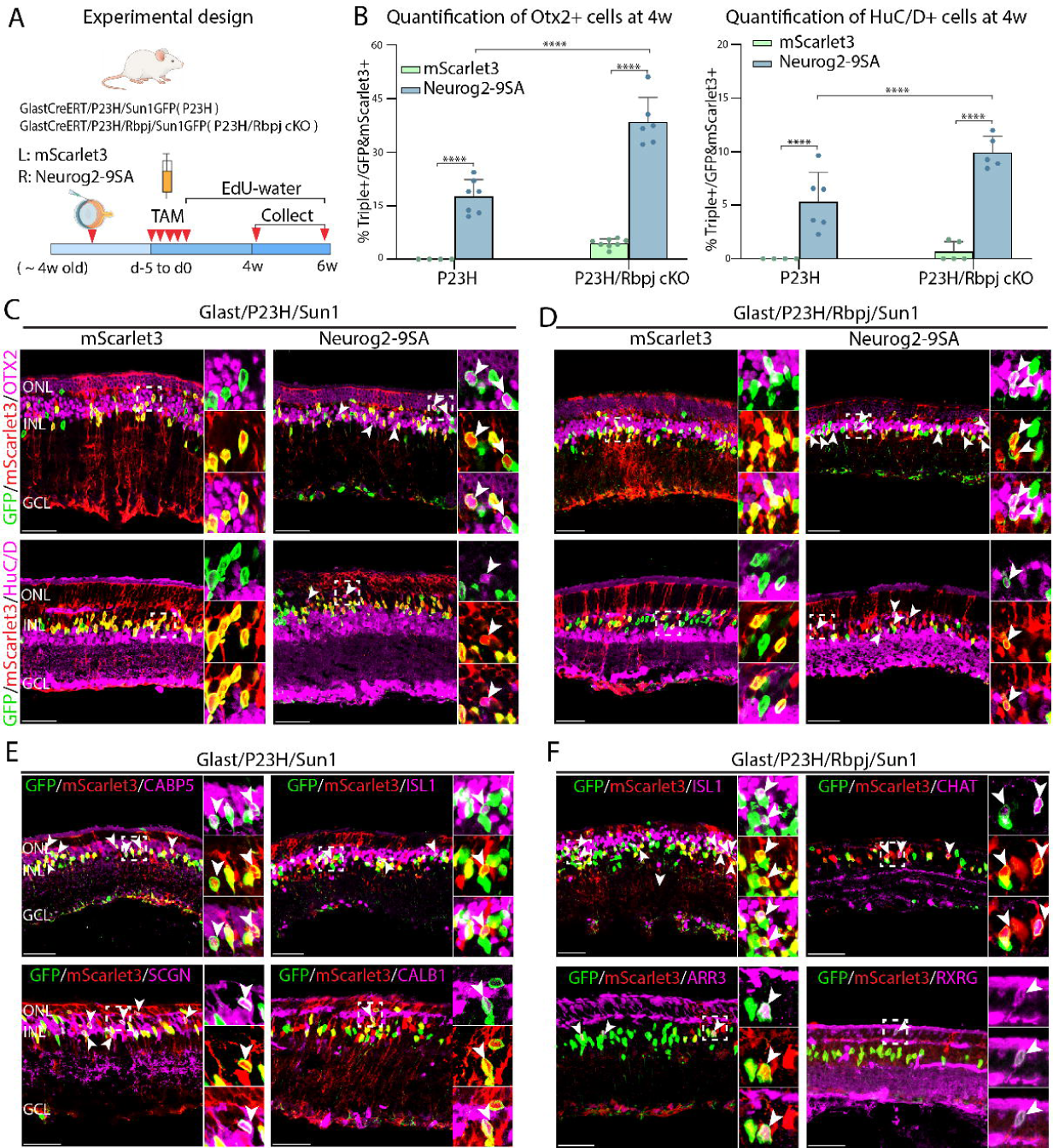
Neurog2-9SA expression reprograms Müller glia to generate retinal neurons in the P23H photoreceptor degeneration model. (A) Experimental design and timeline of sample collection. (B) Quantification of the fraction of GFP⁺/mScarlet3⁺ cells that are OTX2⁺ (left) or HuC/D⁺ (right) in P23H and P23H/Rbpj cKO retinas at 4 weeks post tamoxifen injections. Error bars indicate mean SD. Dot points represent individual retinas. (C, D) Representative images of retinal sections from P23H (C) and P23H/Rbpj cKO (D) mice injected with either mScarlet3 or Neurog2-9SA, immunostained for GFP (Müller glial lineage), mScarlet3 (AAV reporter), and OTX2 or HuC/D. (E) Representative immunostaining images of additional markers, including bipolar/amacrine markers CABP5, ISL1, SCGN and CALB1 in the Neurog2-9SA-injected P23H retinas. (F) Representative images of immunostaining for ISL1, CHAT, and cone markers ARR3 and RXRG in P23H/Rbpj cKO retinas expressing Neurog2-9SA. White arrowheads denote triple-positive GFP⁺/mScarlet3⁺/marker⁺ cells. Müller glia-derived neurons typically exhibit lower GFP intensity than Müller glia.Error bars indicates mean SD. Significance was determined via one-way ANOVA with Tukey’s multiple comparison test: *p < 0.05, **p < 0.01, ***p < 0.001, ****p < 0.0001. ONL: outer nuclei layer; INL: inner nuclear layer; GCL: ganglion cell layers. Scale bars, 50 µm.

We also tested whether inhibition of Notch signalling could enhance neurogenesis by injecting the same AAVs into GlastCreERT/P23H/Rbpj^flox/flox^/Sun1GFP mice (hereafter P23H/Rbpj cKO mice). Neurog2-9SA overexpression resulted in markedly enhanced Müller glia reprogramming, yielding 42% OTX2+ cells and 10% HuC/D+ cells (Fig. 6B, D). These neurons express the cone bipolar cell marker ISL1 and the starburst amacrine marker CHAT (Fig.6F). We found that a subset of generated neurons expressed cone photoreceptor markers ARR3 and RXRG (Fig.6F), as observed in the Neurog2-9SA-infected Rbpj cKO retinas (Fig.5D, Fig.S4A). We did not observe the generation of rod bipolar cells (PRKCA) or RGC (BRN3A and RBPMS), in either P23H or P23H/Rbpj cKO retinas (Fig.S5A, B). EdU tracing data suggests that Neurog2-9SA induces neurogenesis primarily through direct transdifferentiation of Müller glia in both P23H and P23H/Rbpj cKO retinas (Fig.S5C, D).

## Discussion

In this study, we demonstrate that AAV-mediated delivery of a stabilized Neurog2-9SA, is sufficient to reprogram adult Müller glia into multiple types of retinal neurons in the context of both injury-induced and inherited retinal degenerations. Through rigorous lineage tracing, immunohistochemistry, single-cell transcriptomic and epigenomic profiling, we show that Neurog2-9SA activates endogenous neurogenic programs, suppresses Müller glial identity, and drives the formation of neurons that acquire molecular and morphological features of native retinal neurons. Electrophysiological analysis further reveals that these induced neurons exhibit appropriate resting membrane potentials and robust light-evoked responses, indicating successful integration into existing synaptic circuits. Our findings establish Neurog2-9SA as a promising candidate for clinically relevant strategies to induce retinal regeneration from endogenous Müller glia.

Our work expands the current understanding of glia-to-neuron reprogramming in several ways. Previous approaches to reprogramming Müller glia in mammals have often relied on complex transgenic systems and pharmacological treatments, which limit their translational feasibility. For example, our prior studies showed that transgenic deletion of NFIa/b/x transcription factor family ^6^ or Notch signaling ^7,8^ produced primarily retinal interneurons. Similarly, transgenic Ascl1 overexpression often requires retinal injury, histone deacetylase inhibition and yields interneurons ^9^ and ganglion-like cells ^10^. Subsequent work combined AAV-mediated Ascl1 overexpression with acute retinal injury and histone deacetylase inhibition produces limited neurogenesis, an effect that could be enhanced when pairing with Atoh1/Atoh7 overexpression ^33^. In contrast, our study utilized a single AAV-delivered factor that was effective in both acutely injured and chronically degenerating adult retinas. Moreover, earlier AAV-based strategies, such as Ptbp1 knockdown ^11,12^ or NeuroD1 overexpression ^13,14,16^, have since been challenged by the leaky GFP-driven AAV expression in endogenous neurons, highlighting the need for more rigorous and unambiguous approaches ^3,17–19,21,22^. By utilizing Cre-dependent AAV vectors in combination with Sun1GFP reporter lineage tracing, our study provides definitive evidence for bona fide Müller glia–derived neurogenesis.

Neurog2-9SA overexpression broadens the repertoire of neuronal subtypes that can be generated from adult Müller glia. We observed the generation of multiple bipolar subtypes, including Cabp5, Isl1, and Scgn cone bipolar cells, as well as amacrine types. We found that a subset of reprogrammed cells adopted cholinergic starburst amacrine identities, a highly specialized specified neuronal subtype that has not been generated in previous reprogramming studies. We also observed limited Müller glia-derived cells expressing photoreceptor markers Nrl, Arrestin-3 and Rxrg, indicating limited photoreceptor-like differentiation. Furthermore, the reprogramming efficiency and neuronal diversity remained robust in the P23H model of inherited retinal degeneration, demonstrating that Neurog2-9SA-mediated reprogramming is effective across acute injury and chronic diseases.

Our results also highlight the importance of gene dosage and post-translational regulation in shaping the neurogenic potential of reprogramming factors. Whereas wild-type Neurog2 failed to induce substantial Müller glia reprogramming, the stabilized, phospho-insensitive Neurog2-9SA variant exhibited enhanced activity and protein stability. Although our findings do not formally exclude additional mechanisms, such as increased DNA-binding affinity or improved recruitment of chromatin-remodeling complexes, they strongly suggest that kinase-mediated regulation serves to maintain Müller glia identity, a key barrier to glial reprogramming. This idea is further supported by a recent study showing that inhibition of MAK4Ks activity induces proliferation in Müller glia in a YAP-dependent manner ^34^. Our findings indicate that stabilizing proneural transcription factors may represent a generalizable strategy to enhance reprogramming efficiency.

Single-cell multiomic analysis reveals that Neurog2-9SA initiates a coordinated repression of glial maintenance networks, including Notch signaling and NFI-dependent glial identity programs, while simultaneously activating neurogenic cascades involving Ascl1, Insm1, and members of the NeuroD family. Chromatin accessibility analysis revealed strong enrichment of Neurog2 motifs in open regions, supporting the notion that Neurog2-9SA acts in part as a pioneer transcription factor that remodels the chromatin landscape to enable neuronal gene activation. These results indicate that Neurog2-9SA functions as a direct transcriptional activator to orchestrate the transition from a glial to a neurogenic state.

Our results show that reprogramming occurs predominantly through direct glia-to-neuron conversion. EdU labeling and single-cell omics data indicate minimal Müller glia proliferation during this process. This observation is consistent with previous work showing that Neurog2 promotes cell cycle exit and neuronal differentiation ^25,35,36^. The absence of proliferation limits expansion of progenitor cells, and could potentially deplete the pool of Müller glia if the reprogramming efficiency is further increased. Importantly, limited proliferation can minimize oncogenic risk and support the translational safety of direct fate conversion. Furthermore, we and others have shown that Müller glia proliferation can be stimulated through YAP activation ^7,37–39^, Cyclin overexpression ^40^ or kinase inhibition ^34^, suggesting that combining Neurog2-9SA with pro-proliferative interventions may be a necessary approach to expand the progenitor pool and improve regenerative outcomes.

We show that inhibition of Notch signaling substantially enhances the reprogramming activity of Neurog2-9SA across uninjured, acutely injured and chronically degenerating retinas.This synergistic relationship is consistent with the role of Notch signalling in maintaining glial identity and repressing neurogenic programs. In recent work, we demonstrated that Notch inhibition alone is sufficient to induce limited Müller glia-derived neurogenesis, an effect that can be further enhanced by deletion of NFI transcription factors ^7^ or AAV-mediated Oct4 overexpression ^8^. These findings suggest that Neurog2-9SA-based therapies may benefit from concurrent inhibition of Notch signaling to more effectively unlock neurogenic potential of adult Müller glia. Together, our data demonstrate Neurog2-9SA as a potent factor for Müller glia reprogramming, and show that AAV delivery of a single transcription factor can generate diverse, functional neurons in adult retinas under both injury-induced and inherited retinal degenerative conditions. These findings provide a foundation for developing clinically translatable AAV-based regenerative therapies. Future studies aimed at optimizing neuronal subtype specificity, enhancing photoreceptor generation and evaluating restoration of visual function will be essential for advancing Neurog2-9SA-based reprogramming strategies toward therapeutic application.

## Figure legends

**Figure S1. Expression of Neurog2 during retinal development, Müller glia reprogramming, and after AAV delivery** (A) UMAPs showing Neurog2 expression in neurogenic retinal progenitor cells during retinal development in human (Lu et al., 2020) and mouse (Clark et al., 2019), Left panels show cell-type annotations; right panels show Neurog2 mRNA feature plots (color = normalized expression). Neurog2 expression is enriched in neurogenic progenitors and newly born neurons and is low/absent in mature Müller glia. (B) UMAPs showing Neurog2 expression is induced during Müller glia reprogramming in the conditional NFIa/bx/Rbpj knockout retinas (Le et al., 2024). (C) Representative Immunostained images showing Neurog2 (magenta), mScarlet3 (red), and GFP (green) on retinal sections from mice injected with mScarlet3, Neurog2-WT or Neurog2-9SA. Scale bar, 50 µm.

**Figure S2. Additional molecular characterization of neurons derived from Neurog2-9SA-overexpressing Müller glia** (A) UMAP gene expression for Lhx2, Otx2, Elavl4 (HuD) and Slc18a3 (VAChT) across the integrated scRNA-Seq datasets. (B) Heatmap showing top differentially expressed genes across different cell clusters in the scRNA data. (C) Representative images of Neurog2-9SA-overexpressed retinas immunostained for GFP, mScarlet3, and Müller glial marker LHX2. (D) Representative images of Neurog2-9SA-overexpressed retinas for additional markers: NRL (rods), ARR3 (cones), and PRKCA (rod bipolar cells) and RGC markers (ATOH7, BRN3A and RBPMS), co-stained with GFP and mScarlet3. White arrowheads mark GFP⁺/mScarlet3⁺/marker⁺ cells. MG-derived neuron-like cells often show lower GFP intensity than Müller glia (E) Representative images of EdU incorporation assay showing proliferating cells in mScarlet3- and Neurog2-9SA-overexpressed retinas. Note that the majority of EdU+ cells are not GFP+, likely representing immune cells. Scale bars, 50 µm.

**Figure S3. Additional analysis of scMultiomic data.** (A) UMAP plots showing expression of mScarlet3 and Neurog2-9SA mRNA in the mScarlet3 control and Neurog2-9SA samples. (B) Differentially expressed genes across the cell types. (C) Heatmap showing differentially expressed genes along the pseudotime from MG to MGPC. MG, Müller glia, MGPC; Müller glia-derived progenitors; BC, bipolar cell; AC, amacrine cell.

**Figure S4. Additional molecular characterization of neurons generated by Neurog2-9SA expression in Rbpj cKO mice** (A,B) Representative images of Rbpj cKO retinas immunolabeled for additional neuron markers: bipolar (ISL1 and SCGN), rods (NRL), cone (ARR3, RXRG and S-Opsin), rod bipolar (PRKCA) and RGC (BRN3A and RBPMS), GFP and mScarlet3 in uninjured (A) and injured (B) retinas. White arrowheads mark triple-positive GFP⁺/mScarlet3⁺/neuronal marker⁺ cells. (C) Representative images of EdU labeling with GFP and mScarlet immunostainings in mScarlet3 control and Neurog2-9SA overexpressed retinas after NMDA injury. White arrowheads mark GFP-/EdU+ cells, likely representing immune cells. (D) Quantification of Otx2⁺ and HuC/D⁺ percentage among GFP⁺/mScarlet3⁺ cells across three conditions: Rbpj cKO without injury, and Rbpj cKO with NMDA for eyes injected with mScarlet3 or Neurog2-9SA, at 3 weeks post NMDA damage (4 weeks post tamoxifen injections). Error bars indicates mean SD. Significance was determined via one-way ANOVA with Tukey’s multiple comparison test: *P < 0.05, **p < 0.01, ***p < 0.001, ****p < 0.0001. ONL: outer nuclei layer; INL: inner nuclear layer; GCL: ganglion cell layers. Scale bars, 50 µm.

**Figure S5. Additional molecular characterization of neurons generated by Neurog2-9SA expression in the P23H photoreceptor degeneration model.** (A) Representative images of Neurog2-9SA-infected P23H retinas immunolabeled for amacrine cell marker CHAT, rod bipolar marker PRKCA, rod marker NRL, cone marker ARR3 and RGC markers (BRN3A and RBPMS). White arrowheads mark triple-positive GFP⁺/mScarlet3⁺/marker⁺ cells. GFP labels the Müller glial lineage and mScarlet3 marks AAV-transduced cells. (B) Representative images of Neurog2-9SA-infected P23H/Rbpj cKO retinas stained for RGC markers (BRN3A and RBPMS). Müller glia-derived neuron-like cells often display lower GFP intensity than Müller glia. (C,D) Representative images of EdU assay to detect proliferated cells (white arrowheads) in the P23H retinas (C) and in P23H/Rbpj cKO retinas (D) injected with mScarlet3 control or Neurog2-9SA AAV. White arrowheads mark GFP-/EdU+ cells, likely representing immune cells.Scale bars, 50 µm.

**Table 1.** Differential gene expression analysis among cell clusters in the scRNA-seq of mScarlet3 control and Neurog2-9SA at 5 weeks and 2 months post NMDA injury

**Table 2.** Changes on gene expression, chromatin accessibility and transcription factor motif activity among cell states and cell types in the scMultiomic data of mScarlet3 control and Neurog2-9SA at 1 week post NMDA injury

**Table 3.** Key reagents used in the study

## Methods

### Animals

Mice were housed under a 14-hour light/10-hour dark cycle at the University of Michigan Kellogg Eye Center vivarium, with cage lighting maintained at 300 lux. The *GlastCreER;Sun1-GFP* mice strain used in this study were generated by crossing the *GlastCreER* and *Sun1-GFP* lines developed by J. Nathans at Johns Hopkins. *GlastCreER;Rbpjlox/lox;Sun1-GFP* mice were produced by crossing *GlastCreER;Sun1-GFP* with conditional *Rbpj^lox/lox^*mice (Jackson Laboratory, #034200). *GlastCreER ;P23H;Sun1-GFP* mice were obtained by crossing *GlastCreER;Sun1-GFP* with P23H mice (Jackson Laboratory, #017628). *GlastCreER ;P23H;Rbpjlox/lox; Sun1-GFP* mice were generated by crossing *GlastCreER;Rbpj^lox/lox^;Sun1-GFP* with P23H mice. Mice with P23H heterozygosity were used for experiments due to slower retinal degeneration. All procedures were approved by the Institutional Animal Care & Use Committee (IACUC) at the University of Michigan.

### AAV cloning and production

The CAG promoter drove expression of mouse codon-optimized Neurog2 (WT) or codon-optimized Neurog2_9SA linked to mScarlet3 via a P2A peptide to enable bicistronic expression. Codon-optimized Neurog2_9SA contains 9 serine-to-alanine substitutions (S24A, S66A, S192A, S203A, S205A, S215A, S224A, S231A, S234A). All the coding sequences were synthesized (Genewiz). All vectors were packaged into the AAV2/7m8 serotype by the Boston Children’s Hospital Viral Core. All three AAV constructs were deposited on Addgene (Table 3)

### Intravitreal injection of AAV and NMDA

*GlastCreER;Sun1-GFP*, *GlastCreER;Rbpjlox/lox;Sun1-GFP*, *GlastCreER;P23H;Sun1-GFP*, and *GlastCreER;P23H;Rbpjlox/lox;Sun1-GFP* mice (∼4 weeks old) were anesthetized with isoflurane and injected intravitreally with 1.5–2.0 µL of the indicated FLEX-AAV (∼10 E^13 gc/ml viral stock) using a 33-gauge blunt needle. Four days after AAV delivery, mice received Tamoxifen (Sigma-Aldrich H6278) at 2.0 mg per dose in corn oil (Sigma-Aldrich C8267) per day for 5 consecutive days to activate Sun1GFP and AAV expression. Viral titers for each construct are provided in Table 3. For retinal injury of *GlastCreER;Sun1-GFP* and *GlastCreER;Rbpjlox/lox;Sun1-GFP* mice, 2 µL of 100 mM NMDA in PBS was injected intravitreally using a microsyringe fitted with a 33-gauge blunt needle at 1 week post tamoxifen injections.

### EdU treatment

Mice were treated with EdU in drinking water (0.3 mg/ml, Medchem #HY-118411) from the last dose of Tamoxifen injection until sample collection.

### Immunohistochemistry and imaging

Eyes were fixed in 4% paraformaldehyde (Electron Microscopy Sciences 15710) for 2 h at 4 °C. Retinas were dissected in PBS, cryoprotected in 30% sucrose overnight at 4 °C, embedded in OCT (VWR 95057-838), and cryosectioned at 16 µm. Sections were dried (37 °C, 30 min), washed 3× 5 min in 0.1% Triton X-100/PBS (PBST), and (when applicable) processed for EdU using the Click-iT EdU kit (Thermo Fisher C10340/C10636) following the manufacturer’s instructions. After blocking (10% horse serum, 0.4% Triton X-100 in PBS; 2 h, RT), sections were incubated with primary antibodies (Table 3) overnight at 4 °C. After 3× 5 min PBST washes, sections were incubated with secondary antibodies (Table 3) for 2 h at RT, followed with one wash of PBST, 5 mins in DAPI, 2 more washes of PBST, and mounted in ProLong Gold Antifade (Invitrogen P36935) under coverslips (VWR 48404-453). Images were acquired on a Leica STELLARIS 8 FALCON confocal microscope.

### Cell quantification and statistical analysis

For each retina, two randomly selected sections were analyzed. The percentages of *GFP⁺mSacrlet3⁺OTX2⁺*, *GFP⁺mSacrlet3⁺/HuC/D⁺*, *GFP⁺mScarlet3⁺/CHAT⁺*, and *GFP⁺mSacrlet3⁺/ASCL1⁺* cells were calculated relative to the total number of *GFP⁺mSacrlet3⁺* cells. Each point in bar plots represents an individual retina (biological replicate). Quantifications were performed in Imaris (Oxford Instruments). Graphing and statistical analyses were done in GraphPad Prism 10. One-way ANOVA was used for comparisons across multiple groups; data are presented as mean ± SD.

### Whole-cell recording electrophysiology on retinal slices

Mice were dark-adapted overnight. Under red light, mice were euthanized using CO_2_ followed by cervical dislocation. After enucleation, retinas were isolated and hemisected in room-temperature Ames’ medium bubbled by 95% O_2_ 5% CO_2_, then flattened onto filter discs (0.45-μm HA; Millipore), cut into 250 µm slices using a Stoelting tissue slicer with Feather blades (Ted Pella), kept in room-temperature Ames’ medium and shielded from ambient light for up to 7 hr before recording. Slices were rotated 90°, kept in that orientation by slits in two rows of silicone rubber, positioned under a 60✕ water-immersion objective lens, and superfused by 30 - 32 °C Ames’ medium at 3 - 6 mL s^-1^. GFP and mScarlet were visualized via FITC and rhodamine epifluorescence filter sets respectively, and MG-derived neuron-like cells were selected based on morphology and co-expression of the two fluorescent proteins.

Whole-cell recording electrodes were pulled from borosilicate glass (1.5 mm OD, 0.86 mm ID) on a Narishige PC-10 puller, filled with an internal solution (120 mM K-gluconate, 5 mM NaCl, 4 mM KCl, 2 mM EGTA, 10 mM Hepes, 4 mM Mg-ATP, 7 mM Tris2-phosphocreatine, 0.3 mM Na-GTP, 0.01% Alexa Fluor 568 hydrazide, and KOH to adjust pH to 7.3), had tip resistances of 6 - 9 MΩ, and connected to a Molecular Devices MultiClamp 700B amplifier. All light stimuli were full-field 510 nm light 15.6 log quanta cm^-2^ s^-1^ in intensity, produced by passing the condenser light through a narrowband filter, and delivered to the retinal slice from below the transparent bottom of the superfusion chamber. The intracellular Alexa Fluor 568 fill of each recorded cell was imaged right after recording.

### Retinal cell dissociation and fluorescent activated cell sorting (FACS)

Retinas were dissociated into single cell suspension by following previously described protocol ^41^ with some modifications. Briefly, retinas were dissected in ice-cold Hibernate-A (Gibco A1247501) and dissociated using Papain Dissociation System (LK003150, Worthington). Retinas were incubated in papain enzyme buffer at 4C for 20min, then 30C for 10 min, with tube mixing by inversion every 5 min. Retinas were pelleted by quick centrifugation at 200xg for 3 min, then dissociated in ice-cold HBAG buffer containing Hibernate-A, B-27 supplement (Thermo Fisher 17504044), GlutaMAX (Thermo Fisher 35050061) and DnaseI, by pipetting. Single cell suspension was subjected to gradient centrifugation at 200xg for 7min. Cell pellet was then resuspended in HBAG buffer, filtered through 50um filter and subjected to FACS sorting using BD FACS Discover S8 cell sorter for GFP+ cells in HBAG buffer. To concentrate the cells, cells were mixed with BSA at 0.3% final concentration to reduce potential cell clump before centrifuged at 400xg for 5min, then resuspended in desired volume of HBAG buffer to get 500-1500 cells/ul. Cell count and viability were determined by using 0.4% Trypan blue.

### Single-cell RNA and Multiomic library preparation

For scRNA-Seq, GFP+ cells (∼15k) were loaded into the 10X Genomics Chromium Single Cell System (10× Genomics) and libraries were generated using V4 chemistry following the manufacturer’s instructions. Libraries were sequenced on the Illumina NovaSeq platform (500 million reads per library). Sequencing data were processed through the Cell Ranger 9.0.1 pipeline (10x Genomics) using default parameters

Single-cell Multiomic (snRNA/ATAC) libraries were prepared using GFP+ cells using the 10xGenomic Chromium Next GEM Single Cell Multiome ATAC + Gene Expression kit following the manufacturer’s instructions. Briefly, cells were centrifuged down at 500x*g* for 5 min, lysed in 100 μl of ice-cold 0.1x Lysis Buffer by pipette-mixing four times, and incubated on ice for 6 min total. Nuclei were washed with 0.5 ml of ice-cold Wash Buffer and spun down at 500x*g* for 5 min at 4°C. Nuclei pellets were resuspended in ∼20μl 1XNuclei Buffer and counted using Trypan blue. Resuspended cell nuclei (∼15k) were used for transposition and loaded into the 10X Genomics Chromium Single Cell system. ATAC libraries were amplified with 10 PCR cycles and were sequenced on Illumina NovaSeq with ∼500 million reads per library. RNA libraries were amplified from cDNA with 14 PCR cycles and were sequenced on Illumina NovaSeq at ∼500 million reads per library.

Sequencing data were processed through the Cell Ranger arc 2.0.2 pipeline (10x Genomics) using default parameters.

### Single-Cell RNA-seq analysis

Raw scRNA-seq data were processed using Scanpy ^42^. Cells were filtered to retain those with 800–8,000 detected genes, 1,200–30,000 total counts, and less than 25% of counts mapping to mitochondrial genes. Cells failing these criteria were excluded from downstream analyses. Potential doublets were identified and removed using the Python package DoubletDetection ^43^. Cells with doublet scores above a threshold of 0.8 were excluded from further analysis.

The filtered dataset was normalized, log-transformed, and highly variable genes were selected for downstream analysis. Principal component analysis (PCA) was performed, followed by neighborhood graph construction and clustering using the Leiden algorithm. Clusters were annotated based on the expression of canonical marker genes. Differential expression analysis between experimental groups was performed using the Wilcoxon rank-sum test as implemented in Scanpy. Genes expressed in fewer than 10% of cells in a given group were excluded from the analysis.

### Single-cell Multiomic analysis

Single-cell multi-omics analysis integrating RNA and ATAC data was performed using Seurat (v5.1.0), SeuratObject (v5.0.2), and Signac (v1.13.0) in R. Data were first processed at the individual sample level and then integrated for joint analysis.

Nuclei were filtered to remove low-quality or stressed cells using the following criteria. Nuclei with fewer than 100 ATAC fragments or fewer than 1,000 RNA transcripts were excluded. Cells with fewer than 500 detected RNA features were removed. Cells with a nucleosome signal greater than 2 or with more than 10% mitochondrial reads were excluded.These thresholds ensured that only nuclei with sufficient transcriptional and chromatin accessibility information were retained for downstream analyses.

Filtered samples were combined into a single dataset. Joint dimensionality reduction and clustering were performed on RNA and ATAC modalities to identify distinct cell populations. Clusters were refined through iterative reclustering to resolve subpopulations with subtle differences in chromatin accessibility or transcriptional activity. Marker genes for each cluster were identified and annotated using reference gene annotations from a modified GTF reference genome.

Differential gene expression analysis was performed to identify cluster-specific transcriptional programs. Chromatin accessibility was compared across clusters to detect regions with significantly different accessibility. Differential peaks were annotated using the modified reference genome to assign nearby genes and genomic features. The most highly accessible peaks were visualized to highlight cluster-specific regulatory regions.

## Data availability

ScRNA-Seq and scMultiomic sequencing data is available at Gene Expression Omnibus under accession number GSE309445. Codes used for this study is available at our GitHub repository https://github.com/SherineAwad/Neurog2/tree/master and https://github.com/SherineAwad/Neurog2_Omics/tree/master

## Author contributions

Conceptualization: M.Y, Z.F, and T.H. Methodology: M.Y, Z.F, S.A, K.Y.W and T.H. Investigation: M.Y, Z.F, S.A, A.C, J.D, M.C, K.Y.W and T.H. Formal Analysis: M.Y, Z.F, S.A, A.C, J.D, M.C, K.Y.W and T.H. Supervision: T.H. Funding acquisition: T.H. Writing—original draft: M.Y, Z.F, S.A, K.Y.W and T.H. Writing—review and editing: M.Y, Z.F, S.A, A.C, J.D, M.C, K.Y.W and T.H.

## Acknowledgments

We thank P.Hitchcock, D.W. Kim, M.Nagashima and members of the Thanh Hoang lab for comments on this manuscript. This work was supported by an award from the Department of Defense (HT94252510776) to T.H and K.Y.W, a young investigator grant from Alcon Research Institute to T.H, and NIH P30 Vision Research Core to the Department of Ophthalmology.

## Notes

### Competing Interest Statement

The authors have declared no competing interest.

