## Supplementary Figures for "AAV-mediated generation of neurons in injury-induced and inherited retinal degenerations"

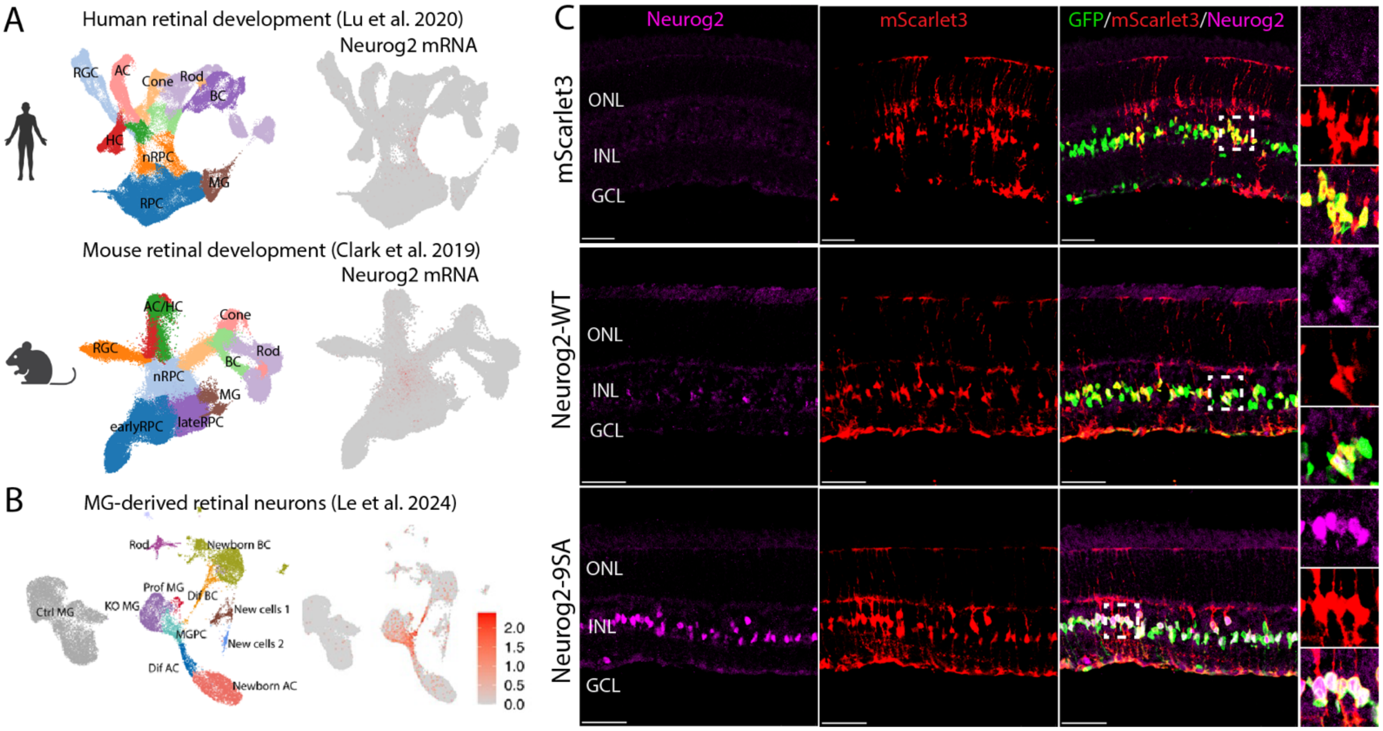


**Figure S1**. **Expression of Neurog2 during retinal development, Müller glia reprogramming, and after AAV delivery.** (A) UMAPs showing Neurog2 expression in neurogenic retinal progenitor cells during retinal development in human (Lu et al., 2020) and mouse (Clark et al., 2019), Left panels show cell-type annotations; right panels show Neurog2 mRNA feature plots (color = normalized expression). Neurog2 expression is enriched in neurogenic progenitors and newly born neurons and is low/absent in mature Müller glia. (B) UMAPs showing Neurog2 expression is induced during Müller glia reprogramming in the conditional NFIa/bx/Rbpj knockout retinas (Le et al., 2024). (C) Representative Immunostained images showing Neurog2 (magenta), mScarlet3 (red), and GFP (green) on retinal sections from mice injected with mScarlet3, Neurog2-WT or Neurog2-9SA. Scale bar, 50 µm.

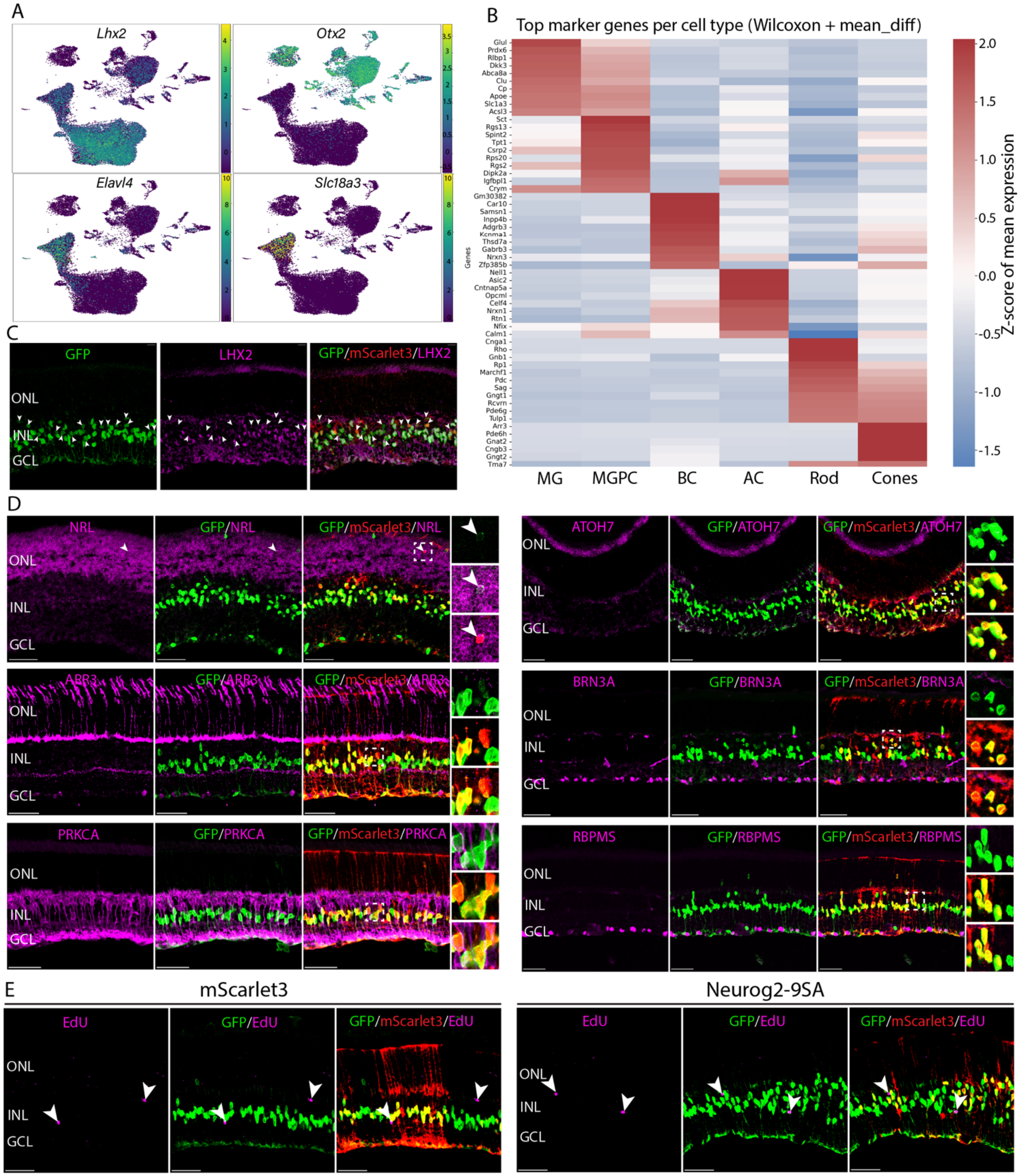


**Figure S2. Additional molecular characterization of neurons derived from Neurog2-9SA-overexpressing Müller glia.** (A) UMAP gene expression for Lhx2, Otx2, Elavl4 (HuD) and Slc18a3 (VAChT) across the integrated scRNA-Seq datasets. (B) Heatmap showing top differentially expressed genes across different cell clusters in the scRNA data. (C) Representative images of Neurog2-9SA-overexpressed retinas immunostained for GFP, mScarlet3, and Müller glial marker LHX2. (D) Representative images of Neurog2-9SA-overexpressed retinas for additional markers: NRL (rods), ARR3 (cones), and PRKCA (rod bipolar cells) and RGC markers (ATOH7, BRN3A and RBPMS) ,co-stained with GFP and mScarlet3. White arrowheads mark GFP$^{+}$/mScarlet3$^{+}$/marker$^{+}$ cells. MG-derived neuron-like cells often show lower GFP intensity than Müller glia (E) Representative images of EdU incorporation assay showing proliferating cells in mScarlet3- and Neurog2-9SA-overexpressed retinas. Note that the majority of EdU+ cells are not GFP+, likely representing immune cells. Scale bars, 50 μm.

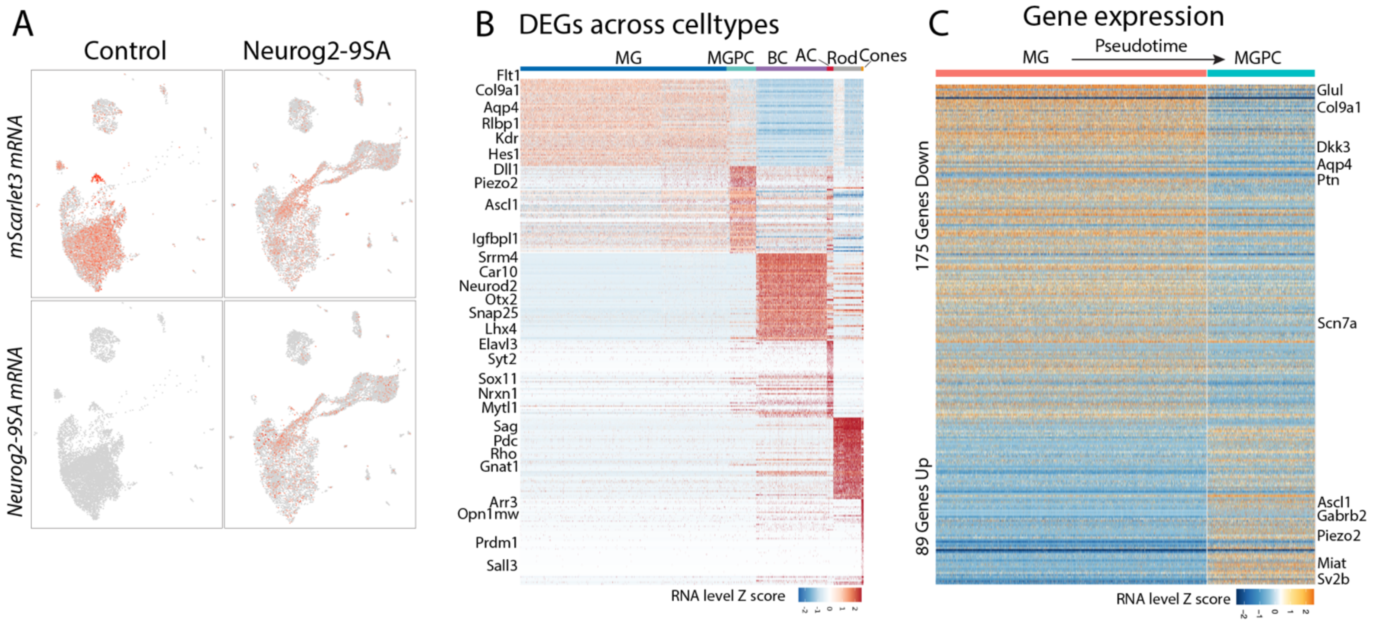


**Figure S3. Additional analysis of scMultiomic data.** (A) UMAP plots showing expression of mScarlet3 and Neurog2-9SA mRNA in the mScarlet3 control and Neurog2-9SA samples. (B) Differentially expressed genes across the cell types. (C) Heatmap showing differentially expressed genes along the pseudotime from MG to MGPC. MG, Müller glia, MGPC; Müller glia-derived progenitors; BC, bipolar cell; AC, amacrine cell.

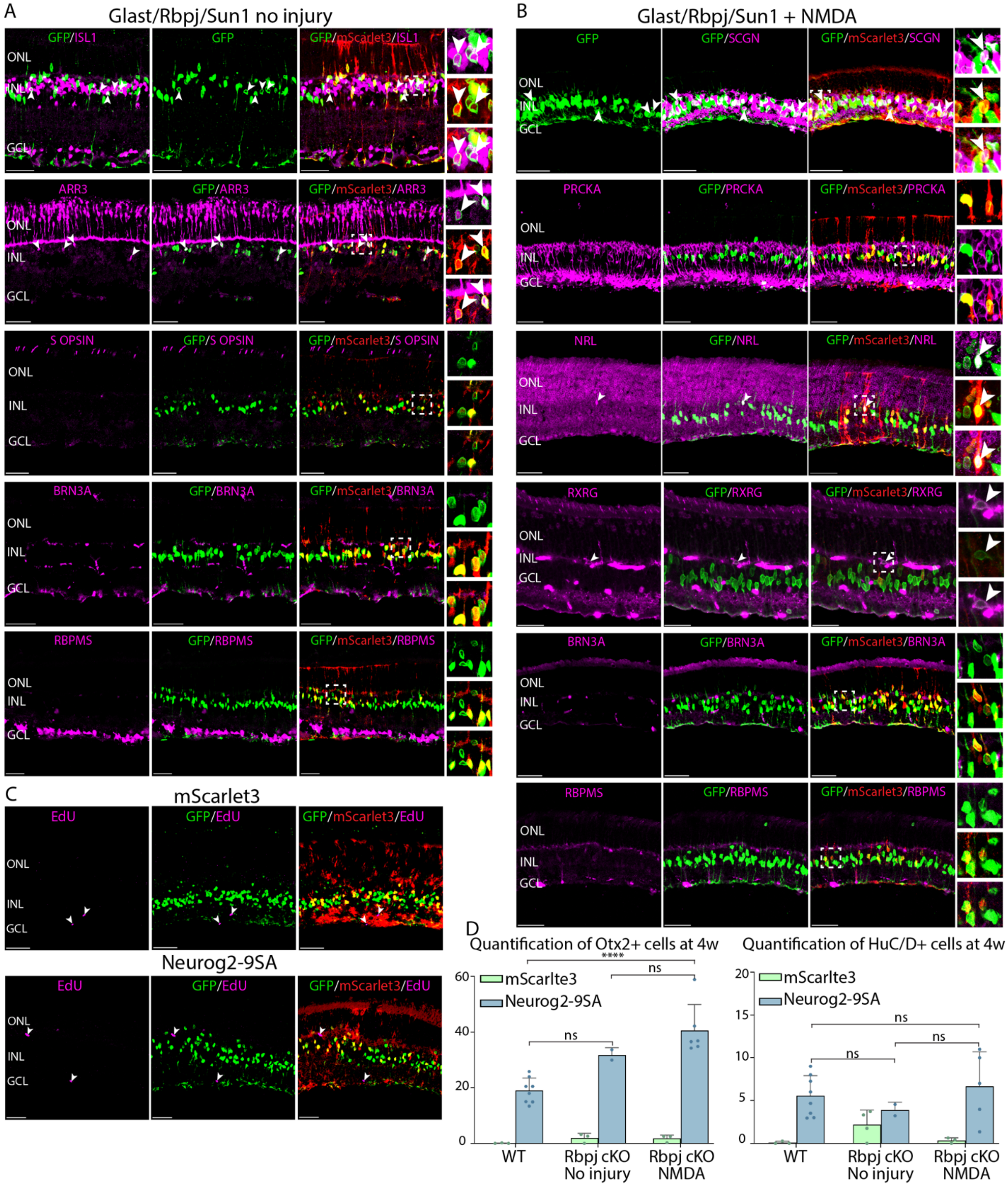


**Figure S4. Additional molecular characterization of neurons generated by Neurog2-9SA expression in Rbpj cKO mice.** (A,B) Representative images of Rbpj cKO retinas immunolabeled for additional neuron markers: bipolar (ISL1 and SCGN), rods (NRL), cone (ARR3, RXRG and S-Opsin), rod bipolar (PRKCA) and RGC (BRN3A and RBPMS), GFP and mScarlet3 in uninjured (A) and injured (B) retinas. White arrowheads mark triple-positive $GFP^+$/$mScarlet3^+$/neuronal $marker^+$ cells. (C) Representative images of EdU labeling with GFP and mScarlet immunostainings in mScarlet3 control and Neurog2-9SA overexpressed retinas after NMDA injury. White arrowheads mark GFP-/EdU+ cells, likely representing immune cells. (D) Quantification of $Otx2^+$ and $HuC/D^+$ percentage among $GFP^+$/$mScarlet3^+$ cells across three conditions: Rbpj cKO without injury, and Rbpj cKO with NMDA for eyes injected with mScarlet3 or Neurog2-9SA, at 3 weeks post NMDA damage (4 weeks post tamoxifen injections). Error bars indicates mean SD. Significance was determined via one-way ANOVA with Tukey's multiple comparison test: *P < 0.05, **p < 0.01, ***p < 0.001, ****p < 0.0001. ONL: outer nuclei layer; INL: inner nuclear layer; GCL: ganglion cell layers. Scale bars, 50 µm.

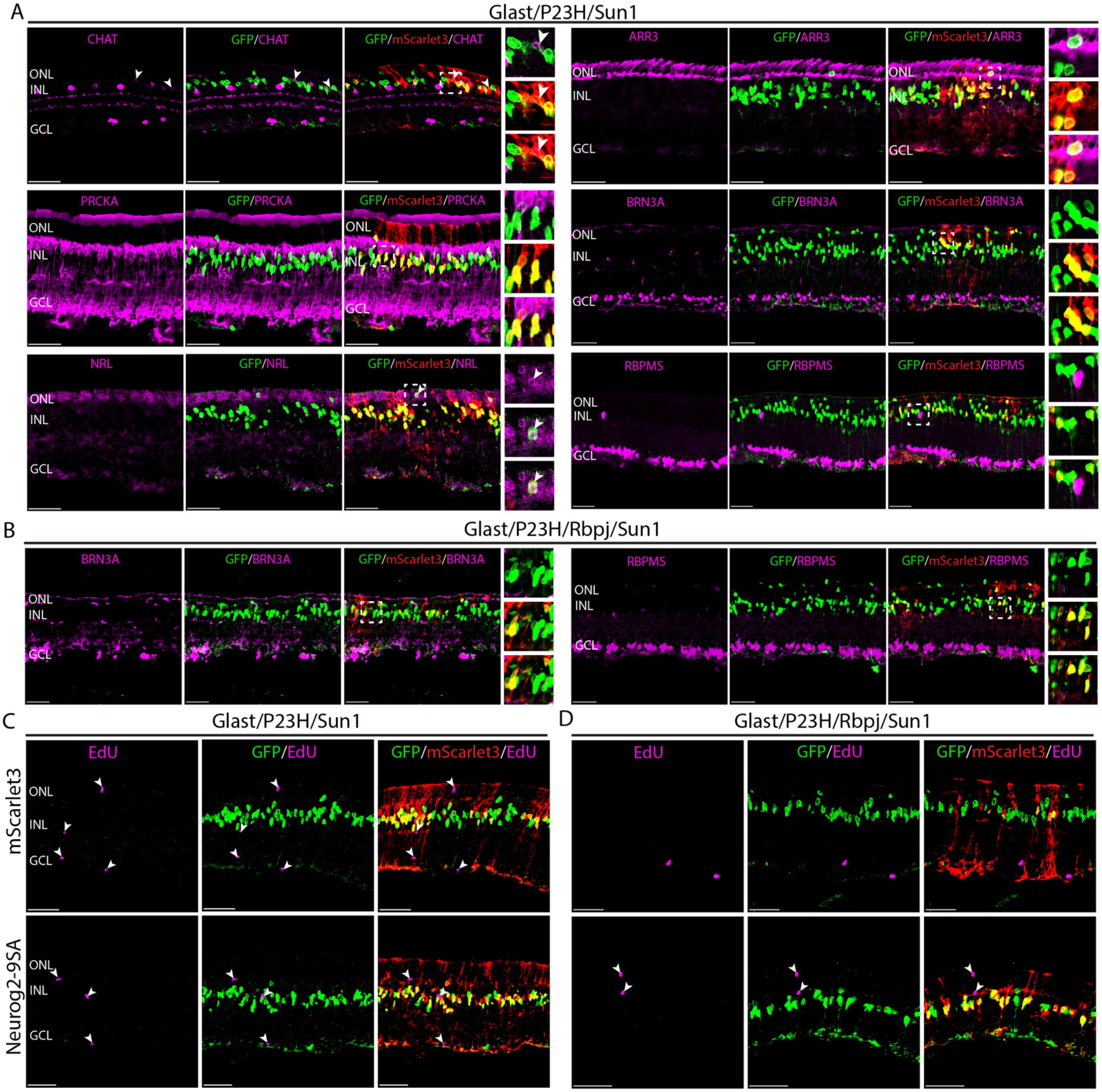


**Figure S5. Additional molecular characterization of neurons generated by Neurog2-9SA expression in the P23H photoreceptor degeneration model.** (A) Representative images of Neurog2-9SA-infected P23H retinas immunolabeled for amacrine cell marker CHAT, rod bipolar marker PRKCA, rod marker NRL, cone marker ARR3 and RGC markers (BRN3A and RBPMS). White arrowheads mark triple-positive GFP$^+$/mScarlet3$^+$/marker$^+$ cells. GFP labels the Müller glial lineage and mScarlet3 marks AAV-transduced cells. (B) Representative images of Neurog2-9SA-infected P23H/Rbpj cKO retinas stained for RGC markers (BRN3A and RBPMS). Müller glia-derived neuron-like cells often display lower GFP intensity than Müller glia. (C,D) Representative images of EdU assay to detect proliferated cells (white arrowheads) in the P23H retinas (C) and in P23H/Rbpj cKO retinas (D) injected with mScarlet3 control or Neurog2-9SA AAV. White arrowheads mark GFP-/EdU+ cells, likely representing immune cells.Scale bars, 50 µm.
